# charisma: An R package to perform reproducible color characterization of digital images for biological studies

**DOI:** 10.1101/2025.11.25.690362

**Authors:** Shawn T. Schwartz, Whitney L.E. Tsai, Elizabeth A. Karan, Mark S. Juhn, Allison J. Shultz, John E. McCormack, Thomas B. Smith, Michael E. Alfaro

## Abstract

Advances in digital imaging and software tools have provided increasingly accessible datasets and methods for analyzing color evolution. Despite the variety of computational packages available, most rely on color classification before running analyses. Previous methods to characterize color limit the ability to analyze large-scale image databases and are not always representative of biologically relevant color classes, which decrease the accuracy of downstream analyses.
Here, we present charisma, an R package designed to characterize the distribution of distinct color classes in images suitable for large-scale studies of biological organisms. Here, we demonstrate the utility of our package through an analysis of color evolution in a sample of diverse and charismatic birds, tanagers, in the subfamily Thraupinae.
We show that charisma can quickly and accurately classify every pixel in an image and validate these results using pre-identified, canonical color swatches. We find that charisma color classifications are consistent with those made by color-pattern experts in the field. Applying charisma to tanager color evolution, we find that charisma outputs seamlessly integrate with downstream evolutionary analyses.
Our results demonstrate that using charisma to manually curate and characterize colors in images provides a standardized, reliable, and reproducible framework for high-throughput color classification.

**Anonymized Data/Code for Peer Review:**

- Analytic data/code for this manuscript can be found on the following Open Science Frame-work repo: https://osf.io/cqg59/overview?view_only=29c9786a338e48618fabbab2883937cc
- charisma package GitHub repo: https://anonymous.4open.science/r/charisma-C87E/

## 1 Introduction

For well over a century, biologists have documented and been intrigued by the charismatic coloration and patterning of earths organisms (Darwin, 1981; Mayr et al., 1963; Santangeli et al., 2023). Animal colors and patterns have been shown to have ecologically important functions such as intra-and inter-specific communication in the form of sexual or social signaling, which includes crypsis, advertisement, or mimicry (Cooney et al., 2019; Feller et al., 2017; Håstad et al., 2005; Irwin, 1994; Rabosky et al., 2016). The evolution of conspicuous coloration and patterning has historically been studied using genetic and observational approaches (Andersson and Amundsen, 1997; Barlow et al., 2018; Ehrlich et al., 1977; McMillan et al., 1999; Neudecker, 1989; Stoddard et al., 2020). However, innovations in digital imaging and novel software tools have provided increasingly accessible datasets and reproducible methods for quantitative ecological and evolutionary investigations of color and pattern (Chan et al., 2019; Endler, 2012; Endler et al., 2018; Hemingson et al., 2024; Maia et al., 2013; Valvo et al., 2020; Van Belleghem et al., 2018; van den Berg et al., 2020; Weller and Westneat, 2019). To date, these approaches have advanced the field’s understanding of the evolution of sexual dichromatism (i.e., males and females differing in color) and the factors influencing why some species have more cryptic, conspicuous, or diverse colors (Cooney et al., 2022; Dale et al., 2015; Shultz and Burns, 2013, 2017; Yu et al., 2024).

Despite this progress, eco-evolutionary explanations for why species display specific colors over others (e.g., blue vs. green) have been lacking. The ability to characterize discrete human-visible color classes in organisms is important to understand the drivers of color evolution and specifically address questions like: Does habitat influence the colors birds evolve? Which colors provide crypsis or conspicuousness? Do specific colors play a role in sexual or social signaling? In birds, color-producing mechanisms are well known for each color; thus, knowing the color classes present in a species can also inform how these mechanisms evolve (Hill and McGraw, 2006; Porzio and Mota, 2025). Given the multidimensional and continuous nature of color and color space, it is both conceptually and technically challenging to carve up the color spectrum into discrete bins, limiting the number of studies and software utilities that use color classification (**Table 1**; Delhey et al., 2023; Ibáñez-Álamo et al., 2025; Nicholson et al., 2007; Senior et al., 2022). Additionally, many color analysis packages require users to input the number of color classes (k) *a priori* to compute various downstream color metrics, which means images must first undergo color profiling such that k meaningfully captures the specific dominant color classes of each organism (**Table 1**; Maia et al., 2019; Van Belleghem et al., 2018; Weller and Westneat, 2019). Previous analyses have been limited to small clades with low color variation across species (Alfaro et al., 2019; Hemingson et al., 2019; Weller and Westneat, 2019). Large-scale color class analyses in birds have used bespoke methods for color classification to analyze bird illustrations (Delhey et al., 2023; Senior et al., 2022). While this allows for broad analyses of color evolution in birds, illustrators exhibit artistic license in their renderings. Color analysis based on images of organisms can reduce these inherent biases. These small datasets and custom methods limit the reproducibility and breadth of organisms that can be studied using this approach.

**Table 1.** Comparison of existing color classification methods.

| Description of color classification method | Considerations | Source | Benefits of charisma |
| --- | --- | --- | --- |
| **Bespoke:** analysis of bird illustrations in CIELab space. A 3D grid of chromatic and achromatic cells was split and merged into 12 color classes. | Bespoke methods are not easily modified or reproduced. Illustrations can be artist biased and may not be available for all organisms. | Delhey et al. (2023);<br>Ibáñez-Álamo et al. (2025) | Flexible and reproducible method that can be used on a variety of image datasets with modifications made simple through the CLUT editor and semi-automated mode. |
| **Bespoke:** analysis of bird illustrations in HSV color space. Hue was divided into 30 degree bins with exceptions for orange, brown, dark, and light for a total of 15 color classes. |  | Senior et al. (2022) |  |
| **Bespoke:** analysis of dewlap pattern, color, and size. Individual color classification from literature and personal images. | Bespoke methods are not easily modified or reproduced. Individual color classifications are not reproducible or feasible for large datasets. | Nicholson et al. (2007) |  |
| **Histogram:** divides color space into regions and computes the proportion of pixels and average pixel value in each region to produce histograms of unique bin centers based on input image. | Single color classes may be broken up across multiple boundaries. | `colordistance`,<br>`recolorize` | The number of potential color classes is user set and the boundaries can be adjusted based on your biological question and system. |
| **K-means clustering:** popular method that partitions pixels into a specified number of bins to minimize difference between the pixel cluster centers. | Fairly slow and biased toward dominant colors. The number of clusters is usually defined *a priori*. | `colordistance`,<br>`patternize`, `pavo`,<br>`recolorize` | Relatively quick, unbiased toward dominant colors, and no a priori k-value necessary. The resulting k-value can be used as input to these other programs. |
| **Receptor Noise Limited Filtering and Clustering:** sequentially groups of segment adjacent pixels according to specific thresholds of spatial and chromatic acuity. | Requires manual designation of thresholds, but can be adjusted. Specific to spatio-chromatic color pattern analysis. | Quantitative Colour<br>Pattern Analysis | General workflow that can be used on its own or integrated into existing color pattern analysis pipelines (e.g. `patternize`, `pavo`). |

To fill this gap, we introduce charisma, an R package designed to streamline the process of characterizing discrete color classes in digital images that can be used in studies of organisms across the tree of life. We provide a flexible and reproducible framework to efficiently determine the presence or absence of key human-visible color classes in images. The charisma package is suitable for large-scale studies of color and color pattern evolution and can be seamlessly integrated into popular downstream quantitative analysis workflows in R (e.g., geiger, patternize, pavo, etc.). We first describe the charisma pipeline and then demonstrate the efficacy of our software by (*i*) validating charisma’s color classification performance using images comprised of predetermined color classes, (*ii*) applying charisma to a small set of standardized, museum specimen images of *Tangara* species and allies in the subfamily Thraupinae and comparing the color classification performance of charisma to those made by an expert, and (*iii*) demonstrating how color classification labels produced by charisma can be leveraged within a trait-based macroevolutionary analysis of color evolution.

## 2 Materials and Methods

### 2.1 The charisma pipeline

The primary function of charisma is to characterize the distribution of human-visible color classes that are present in an image (**Figure 1**). The charisma package was intentionally designed with efficiency and reproducibility in mind, facilitating a standardized and extensible pipeline for color classification. Segmentation of continuous color space into useful classes has previously been done and provides a means to investigate colors and color types in evolutionary biology (Delhey et al., 2023; Ibáñez-Álamo et al., 2025; Senior et al., 2022). To accurately profile the distribution of colors present in an image, we divided hue, saturation, and value (HSV) color space into each of 10 human-visible color classes including all recognized primary and secondary colors (red, orange, yellow, green, blue, and purple) and black, brown, grey, and white (**Figure S1**). We determined HSV color space as the best option for the charisma package for a variety of reasons. (*i*) HSV color space provides an intuitive space to separate human-visible color classes. Using HSV color space makes sense in the context of investigating color class and type evolution regardless of an organism’s visual perception. Incorporating an organism’s visual perception is beyond the scope of this package and requires spectrophotometric measurements or hyperspectral camera imaging and different analyses (Garcia et al., 2015; Hogan and Stoddard, 2024; van den Berg et al., 2020). (*ii*) There is a simple calculation to convert between HSV and red, green, blue (RGB) color spaces, which is useful because most digital images are stored in RGB color space and color distance and complexity are easily calculated in RGB color space. In practice, this conversion can be performed directly using R’s built-in rgb2hsv() function, which returns hue, saturation, and value in normalized units that can then be scaled (e.g., hue × 360*^◦^*, saturation × 100%, value × 100%) for downstream analyses in charisma. (*iii*) An alternative color space is CIELab which was defined by the International Commission on Illumination in 1976 and provides a perceptually uniform color space representative of how the human eye perceives color (Commission Internationale de l’Éclairage (CIE), 2004).

**Fig. 1.**
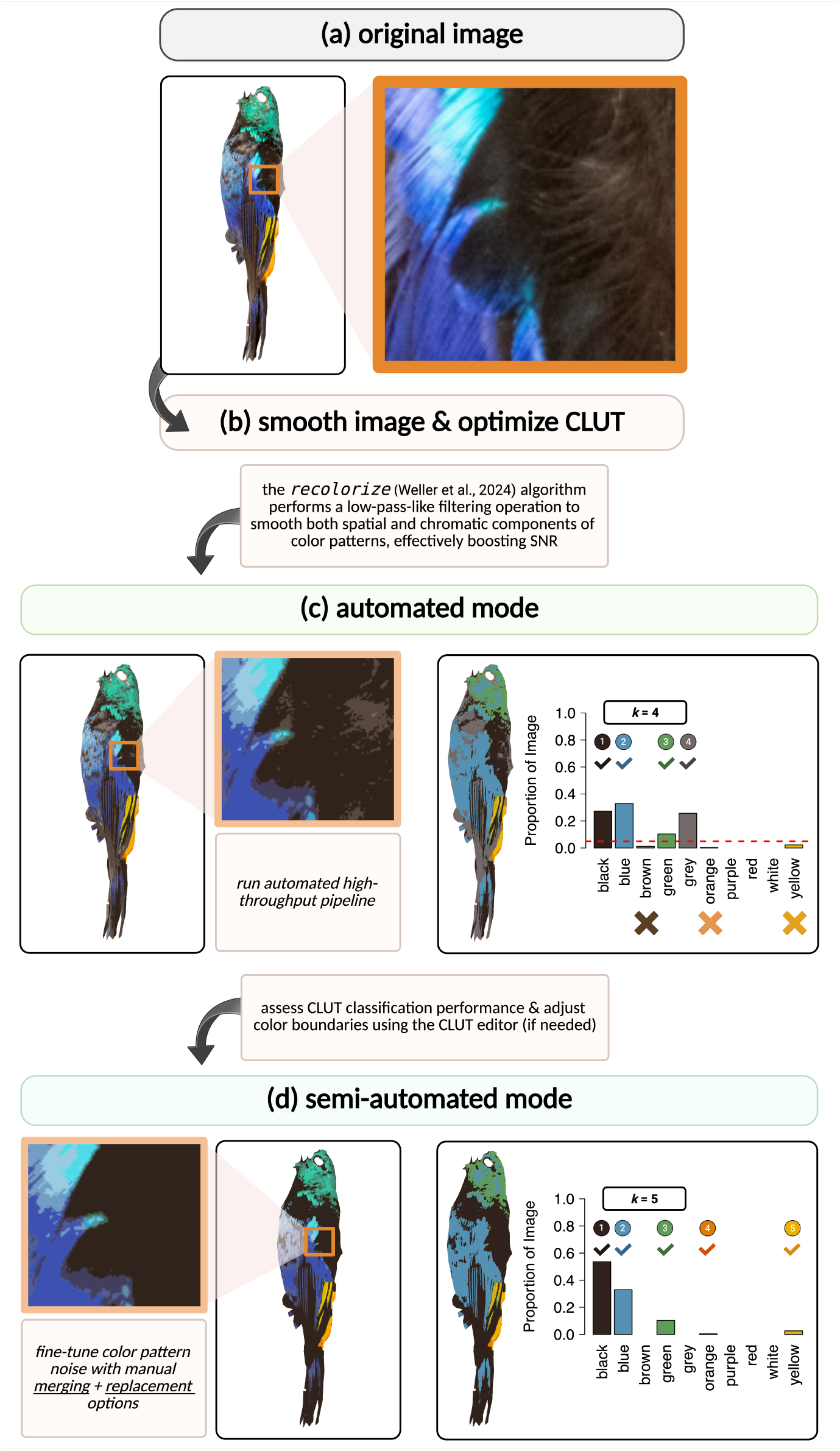
Overview of the *charisma* pipeline. *a,* Original digitized specimen image, *Tangara fastuosa*, passed into *charisma* for processing. *b,* The *recolorize* algorithm (see Weller et al., 2024) is used to boost SNR by effectively performing a low-pass-like filter. *c,* Users run the automated mode to check CLUT performance and adjust the color boundaries according to the image dataset and biological questions using the CLUT editor. *d,* Lastly, users are prompted through the semi-automated mode to make optional, manual adjustments to further fine-tune image noise (e.g., biologically irrelevant colors) with interactive color merging and replacement operations (adapted from the recolorize R package, see above). Users receive the *charisma* “color profile” (right panel in *c* and *d*), which provides a count of the number of discrete color class labels (out of the 10 shown here; *k*) as well as the proportion of pixels in the image that fall into each of the 10 possible color classes. On the left, the dashed red horizontal line indicates a user-specified threshold (5% here) to perform automated rejection of low frequency color classes. In this example, the automated workflow yielded *k* = 4 (i.e., black, blue, green, and grey). The semi-automated mode further reduced noise to eliminate the brown and grey artifacts and correctly call orange and yellow with a 0% threshold.

However, working in CIELab space is biased towards human visual perception, computationally intensive and requires information about lighting conditions, which limits its utility for large scale analyses of images taken in varying light environments or from different sources.

#### Box 1.

##### Recommended workflow for charisma end-users

###### Step 1: Image dataset generation and pre-processing

a. We highly recommend generating an image dataset by taking standardized images of organisms with the same camera, lighting conditions, and color standard (like the X-Rite ColorChecker). We understand that there are advantages to using pre-existing images for color analyses, in which case, we recommend using images from as few sources as possible and only retaining images that can reasonably be compared. For more complete guides on digital imaging for biological studies of color, see Garcia et al. (2015); Stevens et al. (2007).
b. White-balance standardized images using a grey standard. For non-standardized image datasets, more complex color calibration and image standardization is necessary (Akkaynak et al., 2014; Stevens et al., 2007).
c. Segment images manually or use a pre-existing segmentation toolkit like sashimi (Schwartz and Alfaro, 2021) or PixelMator Pro (https://www.apple.com/pixelmator-pro).

###### Step 2: Optimizing the charisma color lookup table (CLUT)

a. Run images through the automated charisma workflow to assess the CLUT classification performance. If the image dataset is less than 50 images, run all images. If the image dataset is much larger, run a subset of the images that includes the breadth of color variation across the dataset.
b. Assess charisma classification outputs and adjust CLUT color boundaries using the CLUT Editor, if necessary.
c. Repeat automated charisma run and CLUT adjustments until satisfied with CLUT performance.

###### Step 3: Finalize charisma outputs

a. Run all images through the semi-automated charisma workflow and manually adjust images using the merge and replacement options, whenever necessary. When merging and replacing colors, think carefully about your biological system question to ensure accurate rep-of colors.

We created a biologically inspired color lookup table (CLUT), which contains distinct, non-overlapping hue, saturation, and value boundaries for each color class (**Figure S1**). To determine the boundaries for each color across the HSV color space, we iteratively developed a schema for defining non-overlapping ranges of hue, saturation, and value triplets for each of the 10 color labels. First, we divided hue into 30 degree bins corresponding to known primary, secondary, and tertiary colors (similar to Senior et al., 2022). We merged and adjusted bins to fit our six hue driven color classes (red, orange, yellow, green, blue, and purple). We defined black, brown, grey, and white on the basis of saturation and value. Then, we used charisma’s automated mode to adjust and fine-tune the color boundaries for our standardized, color-calibrated, research quality photos of bird museum specimens. The CLUT shipped with charisma out-of-the-box is open-source and editable (see charisma::CLUT). It can be validated using the command charisma::validate() to ensure that no color class boundaries are overlapping and that the entire color spectrum is covered. The interactive CLUT editor app, available on the documentation website (https://charisma.shawnschwartz.com/app), allows for simple optimization of the CLUT by adjusting color boundaries and adding or subtracting color classes.

The CLUT defines color boundaries such that a given HSV triplet is classified as belonging to a color category if its hue, saturation, and value simultaneously fall within at least one of the color category’s defined range sets. Moreover, color categories can specify multiple disjunctive range sets (joined by commas), where each set defines a rectangular subregion of HSV space. Within a single range set, the pipe operator (|) signifies a union across non-contiguous intervals within a single dimension (e.g., hue ranges of 0 - 15*^◦^* or 300 - 360*^◦^* for red, which wraps around the hue circle). This syntax enables definition of complex, non-contiguous color category boundaries. For instance, the category “brown” perceptually occupies a non-convex region of color space; therefore, “brown” in the CLUT is defined as the union of several narrow HSV subregions spanning different value and saturation levels. Our interactive CLUT editor provides users the flexibility to define and test their own custom CLUTs (e.g., optimizing classification boundaries for different image sets including non-biological applications) for use within charisma.

Before applying the CLUT-based classification in charisma, the recolorize2() function from the R package recolorize (Weller et al., 2024) is used to produce a spatially ‘smoothed’ version of the input image by removing stray or noisy pixels that may negatively influence the downstream charisma color-label classification (for an example, see **Figure 1**). The recolorize package primarily accomplishes this by performing a spatial-color binning procedure using the RGB ‘histogram’ method (Weller et al., 2024; Weller and Westneat, 2019), which bins each image pixel into a fixed number of color bins. This down-sampled representation of the dominant image colors enables faster color classification by passing each bin’s RGB triplet (i.e., the average color of all pixels assigned to each color bin) into charisma::color2label(r, g, b). The charisma::color2label(r, g, b) function tests for the union of the supplied RGB triplet within the non-overlapping HSV ranges for each color class in the CLUT and returns the matching color label from the set of 10 color classes. This approach is computationally efficient because only a fixed number of bins need to be classified (i.e., 4 bins for each of 3 channels; 4^3^ = 64 cluster centers using the histogram method; cluster centers with a Euclidean distance less than a default cutoff of 20 are combined). The alternative approach classifies every single pixel within the image, which could incorrectly classify shadows or image artifacts and becomes a computationally intensive procedure as the size and resolution of the image scales.

For best results, we recommend using charisma with a multi-stage approach, specifically running both automated and semi-automated workflows together to facilitate accurate color classification fine-tuned for any given set of specimens (Figure 1 and **Box 1**). The automated mode allows users to quickly run all or a subset of images in their dataset to assess the performance of the default CLUT. If users are not satisfied with the charisma ouputs, they should iteratively use the CLUT editor app to adjust the color boundaries and check the results in automated mode. Then, users can use the semi-automated mode to manually merge colors, replace colors, and/or use a threshold cutoff to remove colors with low proportions of pixels that fall below the specified threshold. We developed a custom implementation that draws on the recolorize binning procedure with interactive prompts for color replacement and merging (Weller et al., 2024), empowering users to check the color classification at each stage of the charisma pipeline in an iterative and flexible fashion. The final charisma output (**Figure 1c,d**) for each image includes the number of colors present (*k* = 1…10), a table with presence and absence data for each color class, a log of all manual interventions performed, and R objects that can easily be passed through existing evolutionary analysis pipelines, like those in the R package pavo (Maia et al., 2013, 2019) or analyses of evolutionary models and rates (e.g., Harmon et al., 2008; Pennell et al., 2014; Revell, 2012, 2024). Overall, charisma facilitates a highly standardized and reproducible pipeline to characterize color class data from images of organisms for downstream analyses. By carefully designing the package to handle a spectrum of fully automated to semi-automated workflows, users are equipped to adapt the precision of charisma-derived color label classifications to their particular research needs while enjoying streamlined data and image processing provenance out-of-the-box.

### 2.2 Validating charisma’s color classification performance

We validated the performance of charisma and the accuracy of our CLUT by first testing known color datasets. We obtained color grids for each of our 10 colors from the images on the Wikipedia “Shades of [Color]” pages (**Table S1**). These images contain grids of 9-25 color squares representative of each color, which we used as input in the automated charisma workflow (see **Figure 2**). We also tested images of beetles and fish (**Figure S3**) using the automated mode to provide examples of images that were not used to tune the CLUT (Alfaro et al., 2019; Karan et al., 2021, 2025; Schwartz and Alfaro, 2021; Weller et al., 2024).

**Fig. 2.**
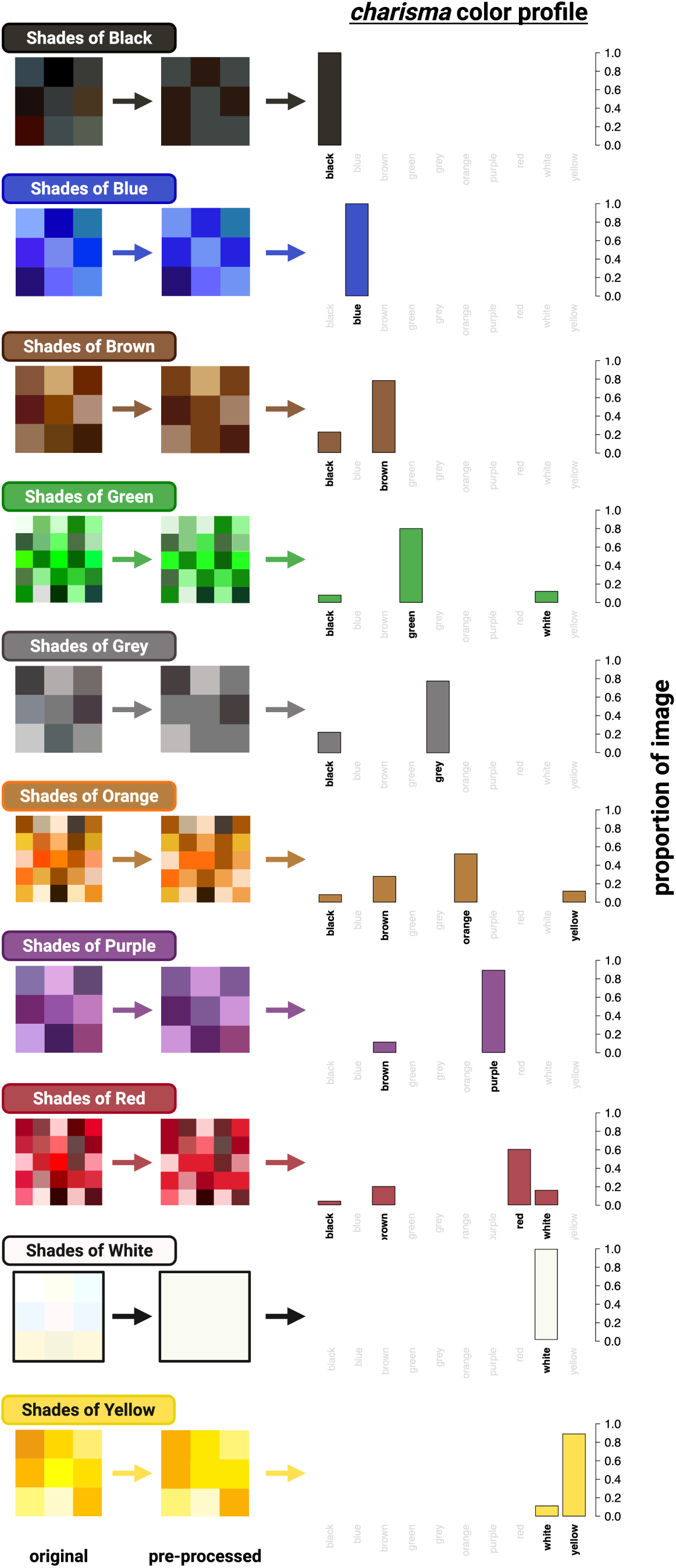
Validating charisma color classifications. Swatches each containing shades of one of ten color classes were obtained from Wikipedia (see **Table S1**) and were then submitted directly to the charisma automated workflow. Charisma performed well-above chance at identifying each of the ten color classes (see *charisma color profile*).

### 2.3 Imaging bird museum specimens

We then illustrate the utility of charisma for evolutionary color analyses with images of tanagers in the subfamily Thraupinae (Family Thraupidae). Tanagers in this subfamily have been well studied in terms of molecular phylogenetics and color evolution (Burns et al., 2014, 2016; Price-Waldman et al., 2020; Shultz and Burns, 2017). Previous work has found that lineages within Thraupinae have the highest evolutionary rates of plumage complexity in tanagers (Price-Waldman et al., 2020). The subfamily Thraupinae also contains the notably colorful genus *Tangara* (and allied genera that have been split from the traditional *Tangara* genus), an ideal clade for testing charisma. We photographed 32 bird museum specimens at the Natural History Museum of Los Angeles County (**Table 4**). This testing dataset consists of incomplete sampling within the subfamily and contains mostly male specimens with a few female specimens that exhibit minimal sexual dichromatism in the human visible spectrum. Specimens were photographed under consistent conditions using a Nikon D70s with Novoflex 35mm lens and Natural LightingNaturesSunlite 30-W full spectrum fluorescent bulbs. Each image included an X-Rite ColorChecker standard and we used the 18% neutral grey standard to white-balance the RAW image files in Photoshop before charisma processing (**Figure S2**). The images were manually segmented by annotating the pixel coordinates to create precise polygonal mask contours directly around the boundary of each bird body (à la Schwartz and Alfaro, 2021) to remove the background. To focus our analyses on plumage coloration, we removed the bill, leg, tag, and cotton eye pixels of the specimen images prior to characterizing color metrics using charisma.

### 2.4 Bird coloration datasets

To classify color in our bird museum specimen images, we ran both the fully automated charisma workflow followed by the semi-automated workflow to manually reduce pixel color noise (**Figure 1**). For the fully automated datasets, we tested threshold values of 5%, 7.5%, and 10%, where any color with a proportion of pixels lower than these threshold values would be removed from the color classification (see **Figure 3**). We implemented these thresholds to automatically remove colors that may have been misclassified due to image artifacts like shadows or feather overlap. For the semi-automated dataset, we merged and/or replaced colors to further refine color clusters for more accurate downstream color classification. To ground-truth our charisma classifications, we extracted color classifications for our species from a dataset of color classifications for all members of the Thraupidae (true tanager) family determined by an expert in tanager coloration (A.J.S.). charisma color classification performance was evaluated by comparing the classifications derived from the automated and semi-automated color datasets against our expert dataset using a binary contingency table (Powers, 2020). We used our expert color dataset as the true colors of the birds and classified each charisma-identified color classification as a true positive (i.e., ‘hit’), false negative (i.e., ‘miss’), false positive (i.e., ‘false alarm’), or true negative (i.e., ‘correct rejection’).

**Fig. 3.**
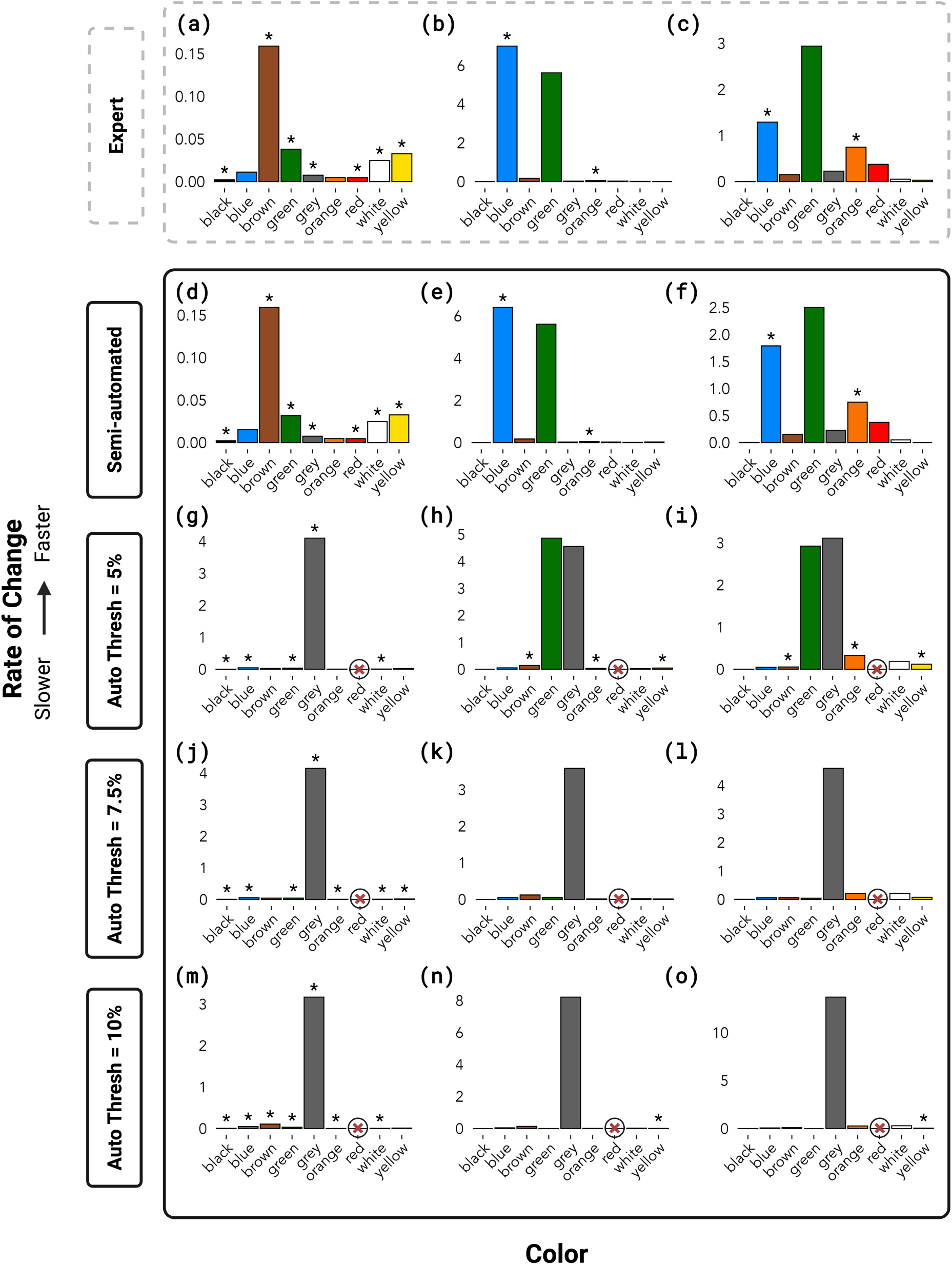
Evolutionary rates of change for each color and evolutionary model as classified in our expert dataset (A.J.S.) – top-dashed – versus charisma-based classifications – Semi-automated: hybrid classification using a threshold = 0% with manual adjustments (W.L.E.T.); the three remaining classifications were derived from the automated charisma workflow with no manual editing, using increasingly conservative color-proportion thresholds from top-to-bottom. Red *⊗* indicates the color was not called by the charisma analysis; asterisks (*) indicate best fit model for each color class (rates and model selection can be found in Table 2). Each plot shows the relative rate of evolutionary change proportional to the highest rate on each plot. Each column represents the following rates: *left,* Equal Rates (ER), *middle,* All Rates Different (ARD) gain, *right,* ARD loss. Note that for the automated dataset with threshold 7.5% (j, k, l), the ER and ARD model selection was inconclusive with an AICc weight of 0.5 for each model.

**Table 2.** Rates of color evolution and ancestral state reconstruction model selection for all datasets.

|  | Color | Equal Rates Model |  |  | All Rates Different Model |  |  |  |
| --- | --- | --- | --- | --- | --- | --- | --- | --- |
|  |  | Rate of gain and loss | AICc | AICc weight | Rate of gain | Rate of loss | AICc | AICc weight |
| **Expert** | Black | 0.00226* | 10.16 | 0.755 | 0 | 0.00226 | 12.41 | 0.245 |
|  | White | 0.0248* | 38.98 | 0.554 | 0.00859 | 0.0505 | 39.42 | 0.446 |
|  | Grey | 0.00749* | 23.75 | 0.565 | 0.0241 | 0.227 | 24.27 | 0.435 |
|  | Brown | 0.159* | 46.21 | 0.75 | 0.169 | 0.152 | 48.41 | 0.25 |
|  | Red | 0.00466* | 18.48 | 0.608 | 0.0252 | 0.374 | 19.36 | 0.392 |
|  | Orange | 0.00475 | 21.4 | 0.266 | 0.0499* | 0.747* | 19.37 | 0.734 |
|  | Yellow | 0.0327* | 41.28 | 0.616 | 0 | 0.0283 | 42.23 | 0.384 |
|  | Green | 0.0380* | 44.87 | 0.589 | 5.61 | 2.94 | 45.6 | 0.411 |
|  | Blue | 0.0111 | 32.8 | 0.42 | 6.99* | 1.29* | 32.15 | 0.58 |
| **Semi-automated** | Black | 0.00226* | 10.16 | 0.755 | 0 | 0.0026 | 12.41 | 0.245 |
|  | White | 0.0248* | 38.98 | 0.554 | 0.00859 | 0.0505 | 39.42 | 0.446 |
|  | Grey | 0.00749* | 23.75 | 0.565 | 0.0241 | 0.227 | 24.27 | 0.435 |
|  | Brown | 0.159* | 46.21 | 0.75 | 0.169 | 0.152 | 48.41 | 0.25 |
|  | Red | 0.00466* | 18.48 | 0.608 | 0.0252 | 0.374 | 19.36 | 0.392 |
|  | Orange | 0.00475 | 21.4 | 0.266 | 0.0499* | 0.747* | 19.37 | 0.734 |
|  | Yellow | 0.0326* | 41.28 | 0.616 | 0.028 | 0 | 42.23 | 0.384 |
|  | Green | 0.0317* | 44.06 | 0.514 | 5.6 | 2.5 | 44.16 | 0.486 |
|  | Blue | 0.0152 | 37.3 | 0.591 | 6.39* | 1.79* | 38.03 | 0.409 |
| **Auto Thresh - 5%** | Black | 0.00226* | 10.16 | 0.755 | 0 | 0.0026 | 12.41 | 0.245 |
|  | White | 0.01* | 27.84 | 0.569 | 0.0278 | 0.188 | 28.4 | 0.431 |
|  | Grey | 4.1* | 46.49 | 0.64 | 4.55 | 3.11 | 47.64 | 0.36 |
|  | Brown | 0.0282 | 42.59 | 0.409 | 0.144* | 0.0563* | 41.85 | 0.591 |
|  | Red | NA | NA | NA | NA | NA | NA | NA |
|  | Orange | 0.00743 | 24.55 | 0.467 | 0.0345* | 0.33* | 24.29 | 0.533 |
|  | Yellow | 0.0243 | 41.845 | 0.486 | 0.0462* | 0.12* | 41.74 | 0.514 |
|  | Green | 0.0359* | 44.87 | 0.72 | 4.86 | 2.92 | 46.75 | 0.28 |
|  | Blue | 0.0472* | 44.92 | 0.747 | 0.055 | 0.046 | 47.09 | 0.253 |
| **Auto Thresh - 7.5%** | Black | 0.00464* | 15.63 | 0.749 | 0 | 0.00463 | 17.82 | 0.251 |
|  | White | 0.00747* | 23.61 | 0.579 | 0.0223 | 0.209 | 24.25 | 0.421 |
|  | Grey | 4.15* | 46.49 | 0.709 | 3.58 | 4.6 | 48.27 | 0.291 |
|  | Brown | 0.0434 | 45 | 0.5 | 0.123 | 0.0648 | 45 | 0.5 |
|  | Red | NA | NA | NA | NA | NA | NA | NA |
|  | Orange | 0.00478* | 18.34 | 0.619 | 0.0141 | 0.205 | 19.31 | 0.381 |
|  | Yellow | 0.0152* | 34.74 | 0.534 | 0.0131 | 0.0769 | 36.02 | 0.466 |
|  | Green | 0.0441* | 45.36 | 0.741 | 0.0582 | 0.0453 | 47.46 | 0.259 |
|  | Blue | 0.0544* | 45.25 | 0.757 | 0.0537 | 0.0567 | 47.51 | 0.243 |
| **Auto Thresh - 10%** | Black | 0.00464* | 15.63 | 0.749 | 0 | 0.00429 | 17.82 | 0.251 |
|  | White | 0.00482* | 18.94 | 0.551 | 0.0198 | 0.292 | 19.35 | 0.449 |
|  | Grey | 3.17* | 46.49 | 0.532 | 8.21 | 13.69 | 46.75 | 0.468 |
|  | Brown | 0.107* | 45.8 | 0.659 | 0.136 | 0.0948 | 47.12 | 0.341 |
|  | Red | NA | NA | NA | NA | NA | NA | NA |
|  | Orange | 0.00229* | 11.83 | 0.675 | 0.00876 | 0.267 | 13.29 | 0.325 |
|  | Yellow | 0.0111 | 29.32 | 0.475 | 0.00748* | 0.0347* | 29.12 | 0.525 |
|  | Green | 0.0317* | 40.99 | 0.63 | 0.00318 | 0 | 42.05 | 0.37 |
|  | Blue | 0.0486* | 44.85 | 0.706 | 0.0484 | 0.0687 | 46.61 | 0.294 |

*Note.* Asterisks (\*) indicate best fit model for each color class.

### 2.5 Evolutionary analyses

We used our datasets and a previously published tanager phylogeny (Burns et al., 2014) to explore variation in the rates of color evolution and reconstruct ancestral color states. Additionally, because the color-producing mechanisms and structures are well known in birds (Hill, 2006; Mason and Bowie, 2020; Porzio and Mota, 2025; Stoddard and Osorio, 2019; Stoddard and Prum, 2011), we also estimated rates and reconstructed ancestral states for color-producing mechanisms. We transformed our color data by grouping discrete colors by avian color-producing mechanism. We designated melanin-based colors as black, brown, and grey, carotenoid-based colors as red, orange, and yellow, and structural colors (produced by feather nanostructure, including those with pigment overlays) as green, blue, and purple (Hill, 2006). For purposes of this study, we have oversimplified the color producing mechanisms for green (produced by carotenoids + nanostructure), and blue and purple (both produced by melanin + nanostructure). We removed white from the color mechanism analysis because it has two potential mechanisms: a lack of pigment or feather nanostructure. For each color and mechanism, we used the fitDiscrete() function in the R package geiger to test the fit of two models of evolution: the equal rates (ER) model, which assumes the rates of gains and losses of a color are equal, and the all rates different (ARD) model, which assumes gains and losses of a color occur at different rates. We compared transition rates for every color in our datasets to determine the potential effects of automated, expert, and semi-automated classifications on evolutionary analyses. We used the sample-size corrected Akaike Information Criterion (AICc) to select the best model for each color and mechanism for our semi-automated dataset. We then leveraged the best-fitting model to reconstruct ancestral states for our semi-automated dataset. Using the R package phytools, we estimated maximum-likelihood probabilities for each color and mapped them on the tanager phylogeny (Burnham et al., 1998; Burns et al., 2014; Huelsenbeck et al., 2003; Paradis et al., 2004; Revell, 2012).

## 3 Results

### 3.1 Validation of charisma’s color classification performance

Figure 2 demonstrates charisma’s ability to characterize the colors in an image. Here, we find that the majority of *charisma* color classifications identify the pre-assigned color class of the input grid. The highest variation in color classifications was present in the red and orange grids with four different colors being called for each. Brown had the second highest proportion of calls for both red and orange grids, which highlights the general difficulty of delineating between the boundaries of these three colors in HSV color space. We demonstrate that diverse sets of organisms can be run using charisma, but we warn users that image lighting and quality can change the designation of the color class depending on the assigned HSV coordinates (see **Figure S3** for examples of beetles and fish images). We stress that users should refer to **Box 1** for our recommended workflow.

### 3.2 Comparison of color datasets

We found that the semi-automated color dataset outperformed automated color datasets when compared to the expert color dataset (**Table 5** and **Tables S2, S3, S4**). Black was well-classified in all datasets and brown and grey had high false alarm rates in automated datasets (**Table 5** and **Tables S2, S3, S4**). The bases and tips of bird feathers, especially contour feathers, are often different colors, with the base generally being black, brown, grey, or white and the tip containing more highly pigmented or structural coloration (Price-Waldman et al., 2025; Terrill and Shultz, 2023). In bird museum specimens, overlapping feathers can be misaligned to create grey or brown patches where the base of feathers show (Figure 1c). These color artifacts may contribute to the over-representation of brown and grey in the automated color datasets. For our analyses, these patches represented unwanted image artifacts, however, depending on the goal of the user, the classification of such subtle variation could be a benefit.

Red and orange had high miss rates in the automated datasets (**Table 5** and **Tables S2, S3, S4**). Red (*n* = 2) and orange (*n* = 1) are rare in our dataset, but when present, they are represented by very small patches. As expected, the automated thresholding procedure removed colors with a small proportion of pixels, making underrepresented color categories difficult to classify using the automated workflow. Together, the small sample size and small patch size of red and orange in these birds contribute to the high miss rates in the automated datasets. The classifications derived from the semi-automated charisma workflow show an almost identical color profile to the expert color dataset as evidenced by the near-100% hit and correct rejection rate (**Table 5**). As such, signal-to-noise ratio (SNR) can be further improved by running the semi-automated mode in charisma to adjust underrepresented color patches (to reduce misses) and manually correct color patches sitting at the boundary of the CLUT color class boundaries (to reduce false alarm identifications).

### 3.3 Evolutionary analyses

To understand how classifying color in different ways (i.e., automated vs. semi-automated datasets) impacts evolutionary rates analyses, we compared evolutionary rates estimated for the expert, semi-automated, and automated datasets (Figure 3 and **Table 2**). We found that the automated datasets elevated the rate of evolution of grey across all models providing evidence that uncorrected color artifacts in bird museum specimens, likely due to misaligned feathers, impact downstream evolutionary analyses. We also found slower evolutionary rates for blue and green in the automated datasets, which may be due to the higher miss rates of charisma-identified blue and green color classes (**Table 5** and **Tables S2, S3, S4**). As predicted, we see high congruence in evolutionary rates between our expert and semi-automated color datasets.

Given the similarity between our expert and semi-automated color datasets, we present the results of our evolutionary analyses using the best-performing charisma classification (i.e., the semi-automated color dataset). We note that these results and interpretations are based on incomplete sampling of a subfamily of tanagers and they are presented as an example of the utility of charisma for these types of evolutionary analyses. Additionally, we have categorized highly complicated color-producing mechanisms into two simple mechanisms. Any biological interpretations of the data should be taken with caution. We tested two models of evolution on discrete color classes and colors grouped by color-producing mechanism (see Figure 4, **Table 2**, and **Table 3**). We excluded purple and the melanin mechanism from our analyses because purple was not present in any species and melanin was ubiquitous across all species. When comparing the evolution of the melanin color using the ER model, brown showed an elevated rate compared to all other color classes, and black showed the slowest rate (Figure 3). Melanin molecules are structurally robust and are often deposited in the primary wing and tail feathers of birds to increase durability and resist abrasion in these high-use feathers (Bonser, 1995). This aligns with our finding that melanin coloration is highly conserved and indicates the need for structural integrity in bird feathers.

**Fig. 4.**
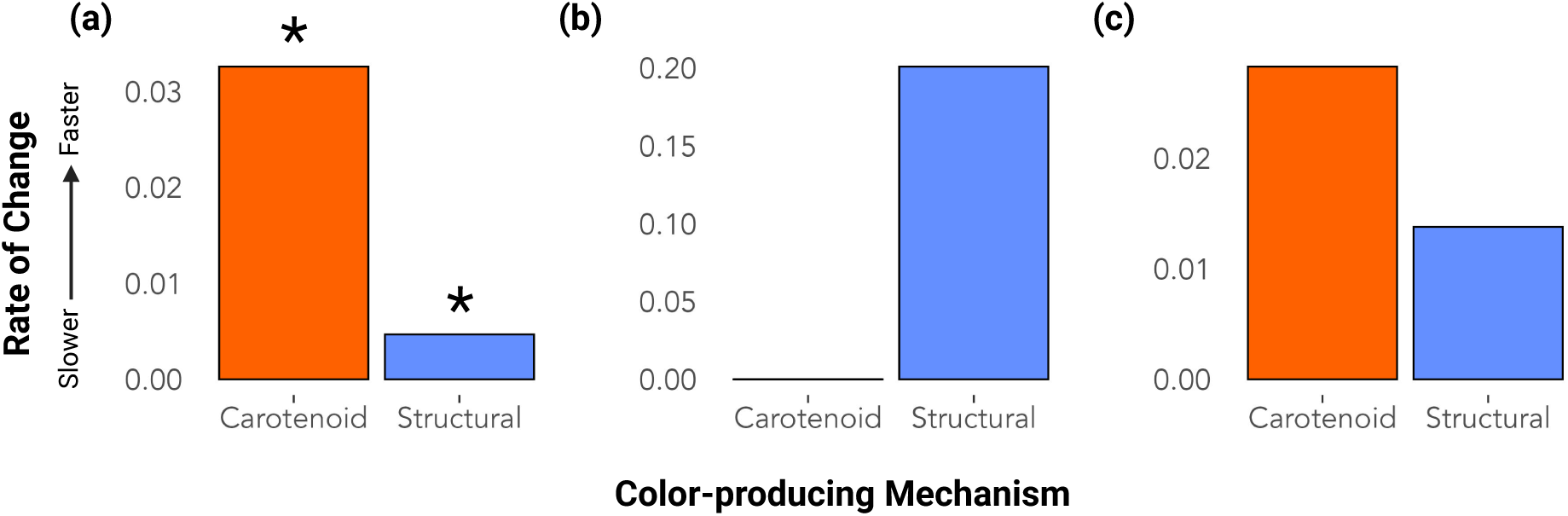
Evolutionary rates for each color-producing mechanism and evolutionary model for the semiautomated dataset: *a,* Equal Rates (ER) rate, *b,* All Rates Different (ARD) gain rate, *c,* ARD loss rate. Asterisks (*) indicate best fit model for each color-producing mechanism (rates and model selection can be found in Table 3).

**Table 3.** Rates of color-producing mechanism evolution and ancestral state reconstruction model selection for the semi-automated dataset.

| Mechanism | Equal Rates Model |  |  | All Rates Different Model |  |  |  |
| --- | --- | --- | --- | --- | --- | --- | --- |
|  | Rate of gain and loss | AICc | AICc weight | Rate of gain | Rate of loss | AICc | AICc weight |
| Carotenoid | 0.0326* | 41.28 | 0.616 | 0 | 0.0283 | 42.23 | 0.384 |
| Structural | 0.00468* | 17.35 | 0.725 | 0.201 | 0.0138 | 19.29 | 0.275 |

*Note.* Asterisks (\*) indicate best fit model for each color class.

**Table 4.**
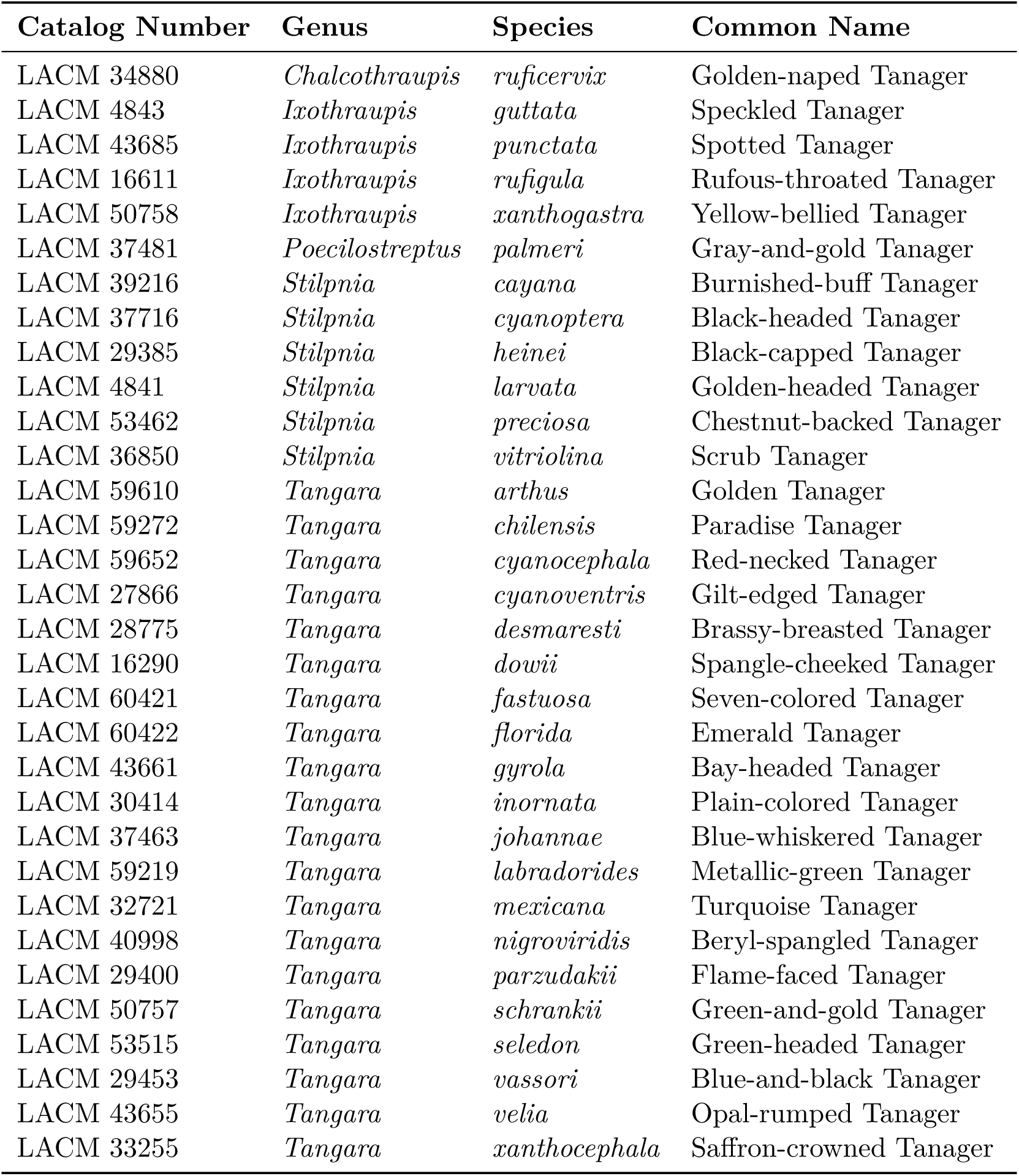
Bird museum specimens.

**Table 5.** Performance comparison between the semi-automated charisma classification workflow (with manual adjustments) and expert classification across color classes. Each column reports the total number of hits (true positives), misses (false negatives), false alarms (false positives), and correct rejections (true negatives) in the charisma dataset. The percentage of correct hits (parentheses) are calculated as the number of hits divided by the total number of color targets (hits + misses) in the expert dataset. The percentage of correct rejections (parentheses) are calculated as the number of correct rejections divided by the total number of non-targets (correct rejections + false alarms) in the expert dataset.

| Color | Hits | Misses | Correct Rejections | False Alarms |
| --- | --- | --- | --- | --- |
| Black | 31 (100%) | 0 | 1 (100%) | 0 |
| Blue | 25 (93%) | 2 | 5 (100%) | 0 |
| Brown | 17 (100%) | 0 | 15 (100%) | 0 |
| Green | 21 (100%) | 0 | 10 (91%) | 1 |
| Grey | 3 (100%) | 0 | 29 (100%) | 0 |
| Orange | 2 (100%) | 0 | 30 (100%) | 0 |
| Purple | 0 (100%) | 0 | 32 (100%) | 0 |
| Red | 2 (100%) | 0 | 30 (100%) | 0 |
| White | 9 (100%) | 0 | 23 (100%) | 0 |
| Yellow | 19 (100%) | 0 | 13 (100%) | 0 |

For color-producing mechanisms, carotenoid coloration showed a higher rate of evolution than structural coloration (Figure 4). Pigment-based color is more widespread across the avian tree of life than structural color (Hill and McGraw, 2006). However, across our tanager dataset, structural colors (green and blue), are present in more species than carotenoid colors (red, orange, and yellow), which may be contributing to the faster rate of evolution for carotenoid coloration. The structural color, blue, best fit the ARD model of evolution and showed elevated gain and loss rates compared to all other colors (Figure 3 and **Table 3**). Where structural color is present, there is evidence that color diversity accumulates faster than pigment-based color because of the modularity of layering pigments and structure (Eliason et al., 2015; Maia et al., 2016). These subtle structural changes at the nanometer level may result in the production of significantly different colors allowing for rapid evolution and accumulation of color diversity.

Lastly, we found the ER model of evolution to be the best-fitting model for every color and color-producing mechanism except for blue and orange (**Table 2**). Using the best-fitting model for each color and mechanism, we reconstructed ancestral states and mapped them across the tanager phylogeny (Figure 5). We found that most ancestral nodes had black, brown, yellow, green, and blue color states, which were also the most common colors in our dataset. Black, white, brown, and grey have been shown to be the most common colors across the bird tree of life (Delhey et al., 2023) and differences in our findings are likely due to the uniquely colorful nature of birds in the subfamily Thraupinae. The ancestral state reconstructions demonstrate that rates of evolution are largely driven by losses of colors across internal branches of the tree. Orange fit the ARD model best with a slightly elevated rate of gain of orange than loss (Figure 3). However, orange is only present in one species in our dataset and this result is driven mainly by the uncertainty of the presence of orange at the root of the phylogeny (see Figures 3 **and 5**).

**Fig. 5.**
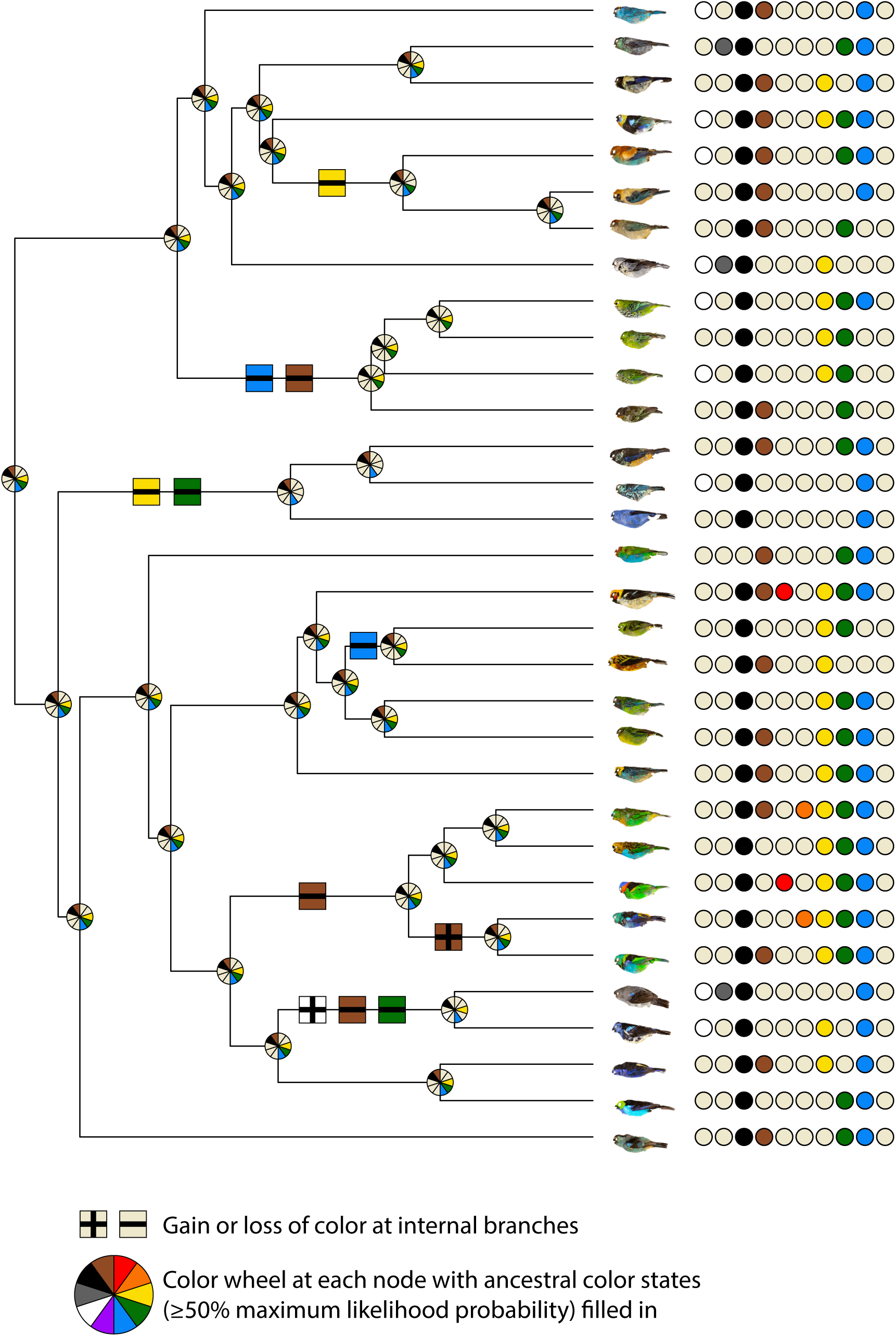
charisma color classifications mapped on the tanager phylogeny. For each species, colored dots represent the colors present in the image. Color wheels at each node indicate estimated colors with greater than or equal to 50% maximum likelihood probability. Internal branches are marked with gains or losses of colors. Species order (top to bottom): *Chalcothraupis ruficervix, Stilpnia heinei, S. cyanoptera, S. larvata, S. preciosa, S. cayana, S. vitriolina, Poecilostreptus palmeri, Ixothraupis guttata, I. xanthogastra, I. punctata, I. rufigula, Tangara dowii, T. nigroviridis, T. vassorii, T. gyrola, T. parzudakii, T. florida, T. arthus, T. johannae, T. schrankii, T. xanthocephala, T. desmaresti, T. cyanoventris, T. cyanocephala, T. fastuosa, T. seledon, T. i1n7ornata, T. mexicana, T. velia, T. chilensis, T. labradorides*.

## 4 Discussion

Here, we present charisma, an R package we developed to provide a standardized, reproducible, and flexible framework for characterizing color classes from digital images. To demonstrate its utility, we used standardized images of bird museum specimens and found that a hybrid approach using both the automated and semi-automated workflows provided the most accurate color profiling. Artifacts from misaligned feathers in our bird museum specimen image dataset led to over representation of brown and grey in the automated charisma-derived color classification. While these artifacts might not generalize to all image datasets, we recommend using charisma in two-stages: first, use the automated mode to check and optimize the CLUT classification performance; then, use the semi-automated mode to adjust the outputs using charisma’s built-in utilities for classification clean-up.

Overall, charisma is built with a fully open source design philosophy where flexibility and customization are both welcomed and encouraged. One example is the built-in functionality to adjust the default CLUT based on user needs and their specific image dataset using the CLUT editor app. Cases where this might be most useful are when fine-tuning is needed for standardized datasets of other organisms, like J.E. Randall’s images of fish (http://pbs.bishopmuseum.org/images/JER/), or for extant datasets comprised of non-standardized images, like those culled from iNaturalist (Matheson, 2014; https://www.inaturalist.org) or eBird (Sullivan et al., 2009). We note that image quality and variation can drastically impact downstream analyses and considering them is critical for accurate analyses of color evolution. We warn users against blindly using the default CLUT without carefully checking color classifications and refer users to our use recommendations in **Box 1**.

Lastly, charisma enables seamless transition to downstream evolutionary analyses, which we show-cased in a vignette on tanagers in the subfamily Thraupinae. More broadly, we present charisma as a solution for high-throughput organismal color analyses with utility that is not restricted to evolutionary biology applications.

## Acknowledgments

We extend our gratitude to current and former members of the Alfaro Lab, Center for Tropical Research, and Moore Laboratory of Zoology whose feedback greatly influenced the development of charisma. Specifically, we thank Mackenzie Perillo and Kevin Wang for their help in preparing the testing and validating datasets for charisma, Gregory Grether for helpful comments on earlier versions of this manuscript, and Hannah Weller for insightful and inspiring conversations about color pattern evolution, computational methods for classifying color, and the development of both the colordistance and recolorize R packages, of which charisma was inspired by and builds upon. Specimens were provided by the Natural History Museum of Los Angeles County.

## Supplementary Information

**Fig. S1.**
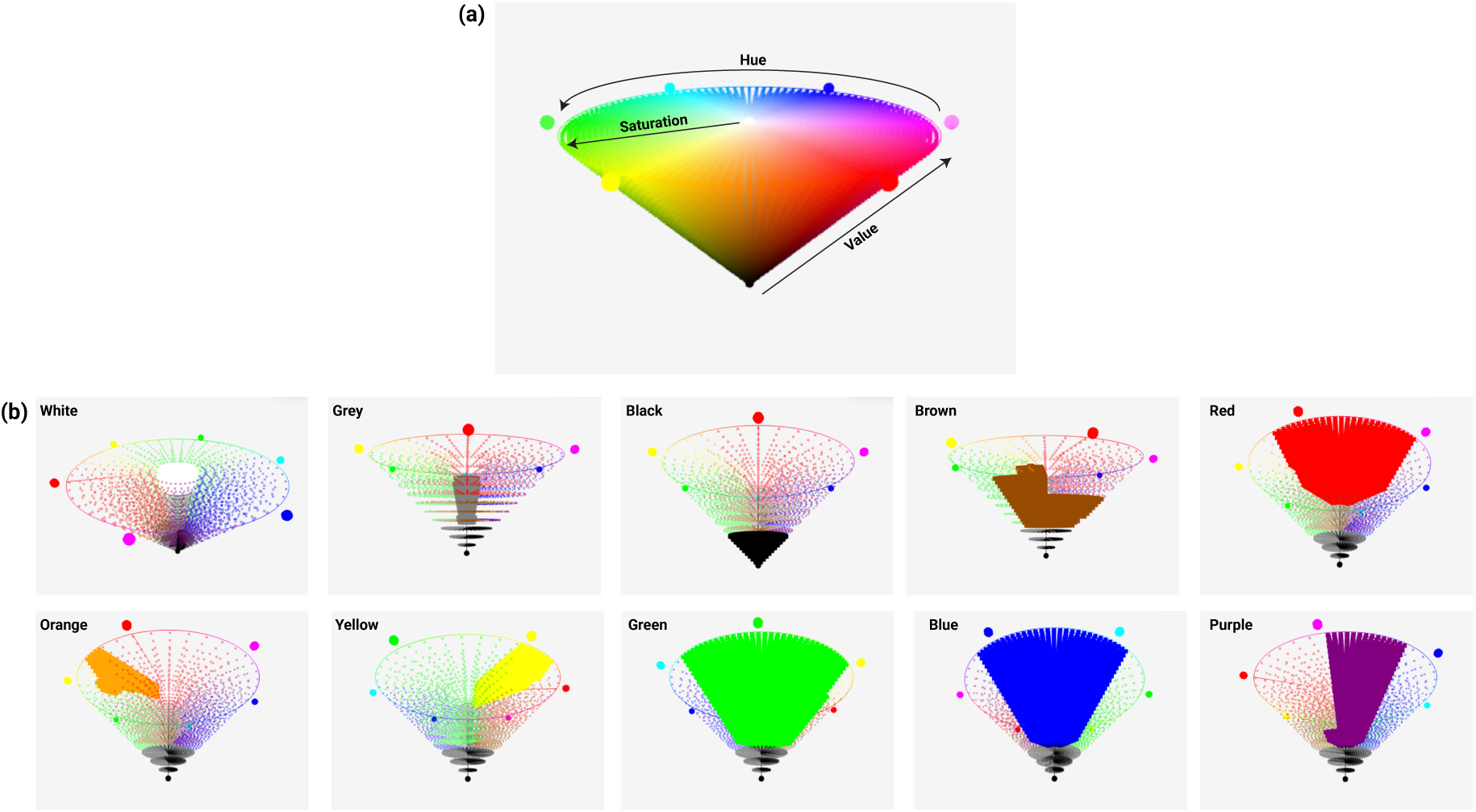
Full HSV color space (a) and color slices for each color class (b) as visualized using the CLUT editor.

**Fig. S2.**
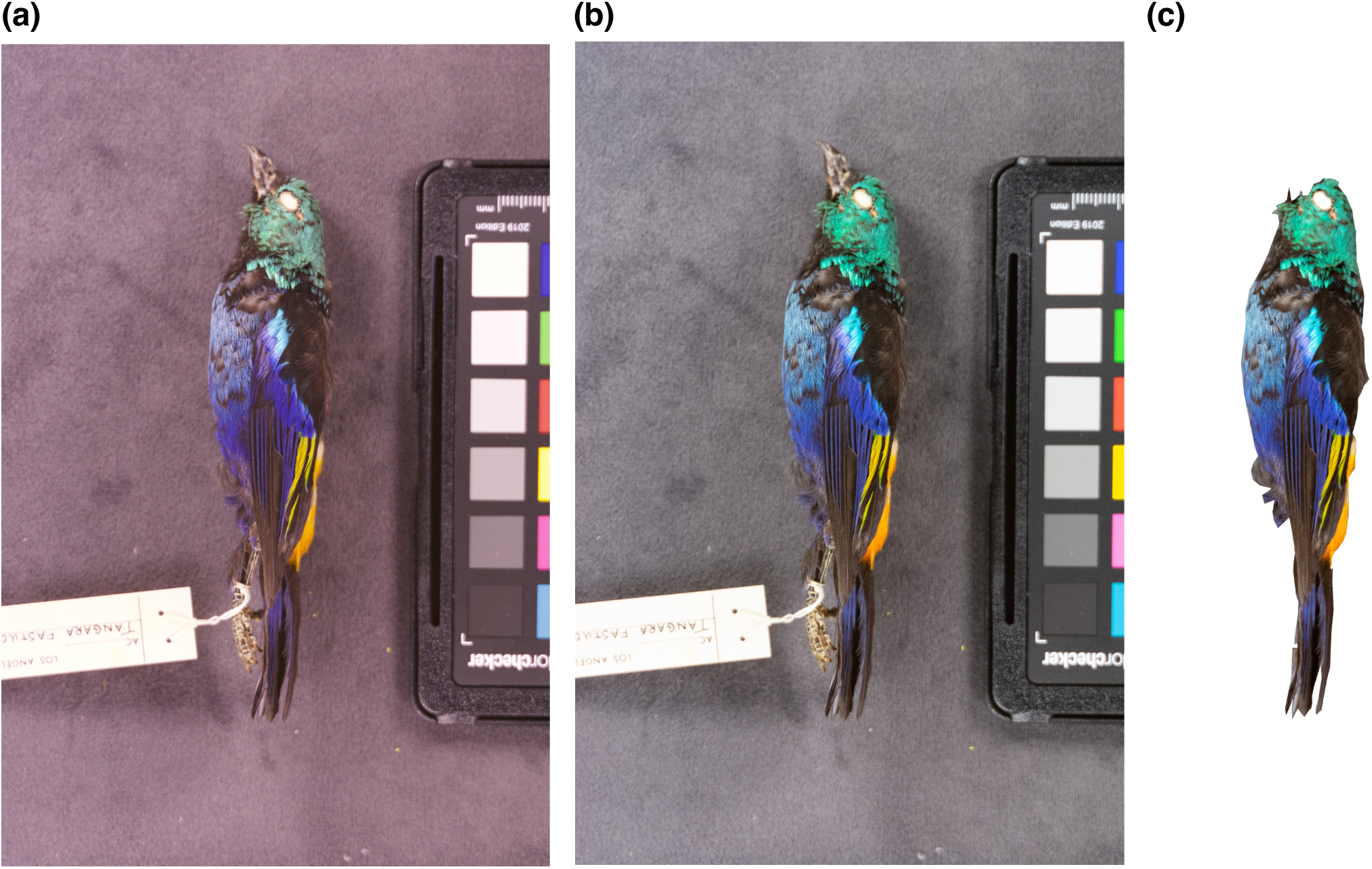
Comparison of original image (a), white-balanced image (b), and white-balanced and segmented image

**Fig. S3.**
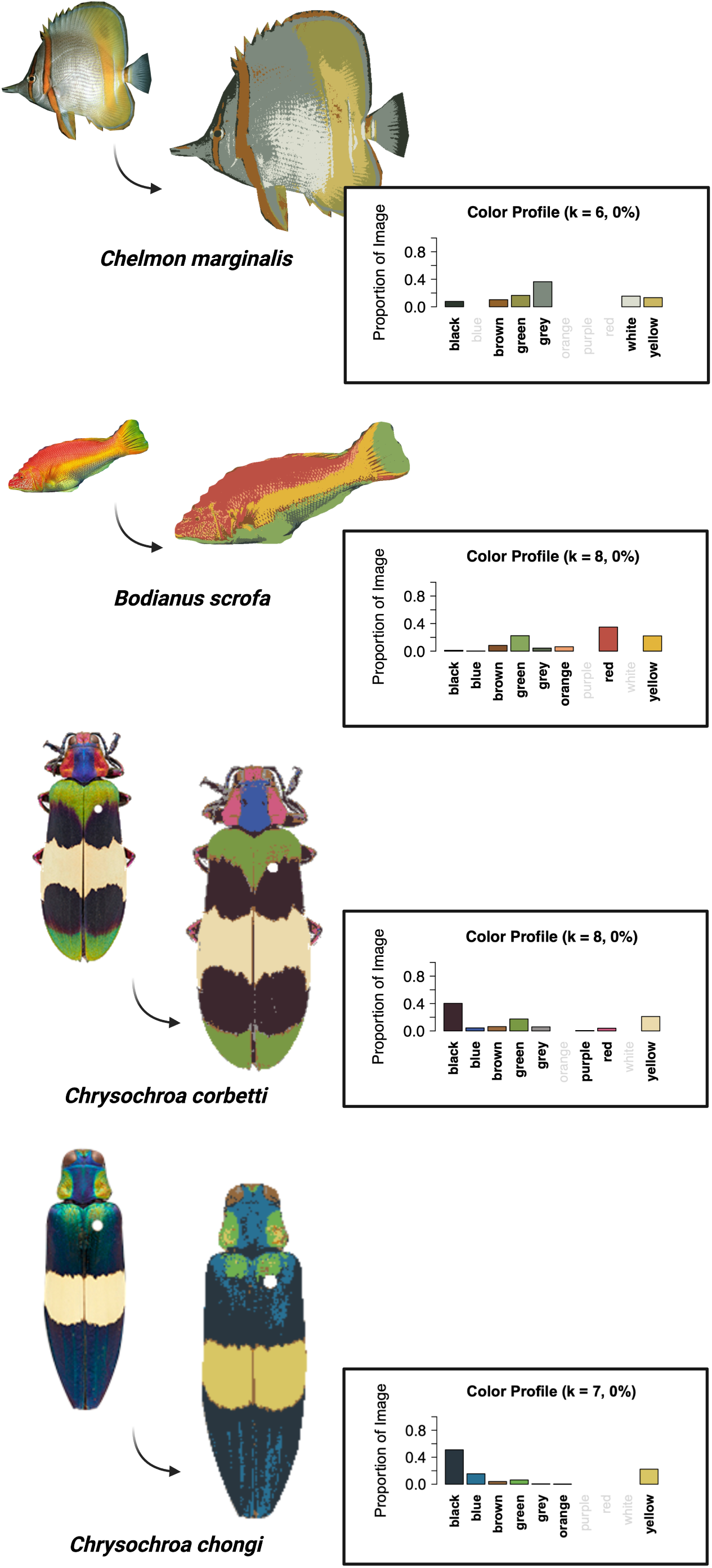
charisma color classifications for fish (Alfaro et al., 2019; Karan et al., 2021, 2025; Schwartz and Alfaro, 2021) and beetles (Weller et al., 2024)

---

**Table S1.**
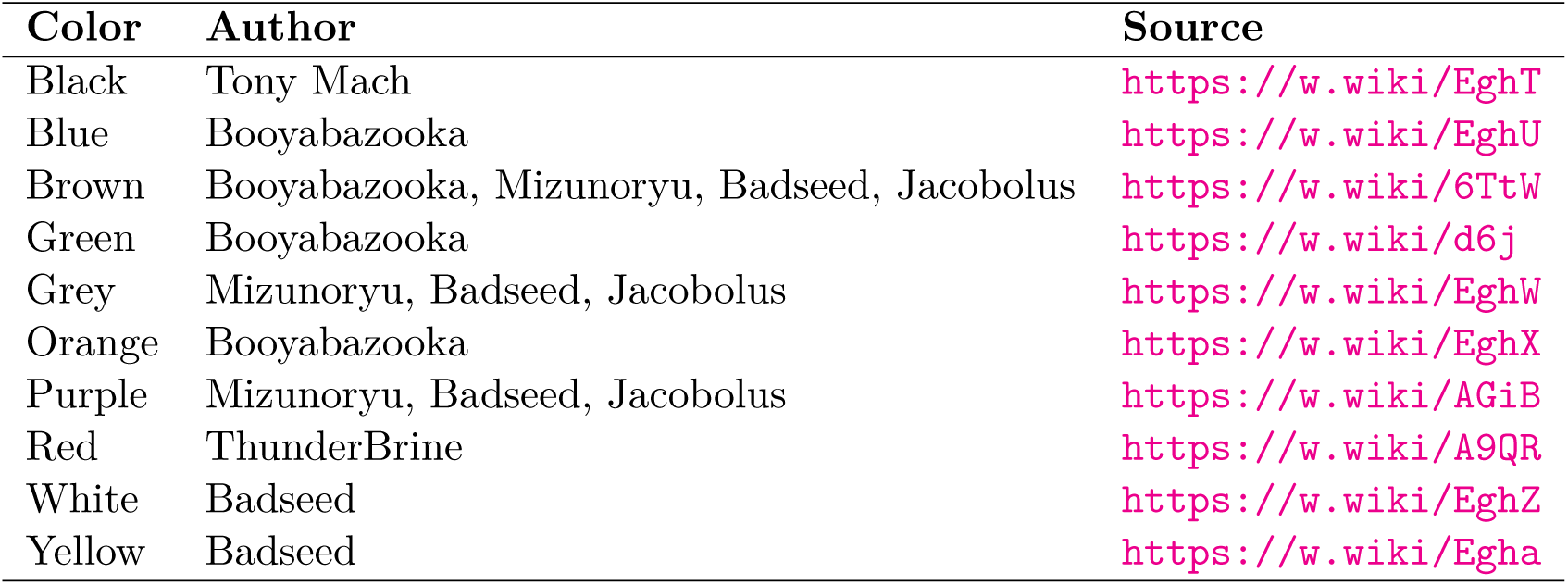
Sources images for Wikipedia-based shades of color swatch validation.

| Color | Author | Source |
| --- | --- | --- |
| Black | Tony Mach | https://w.wiki/EghT |
| Blue | Booyabazooka | https://w.wiki/EghU |
| Brown | Booyabazooka, Mizunoryu, Badseed, Jacobolus | https://w.wiki/6TtW |
| Green | Booyabazooka | https://w.wiki/d6j |
| Grey | Mizunoryu, Badseed, Jacobolus | https://w.wiki/EghW |
| Orange | Booyabazooka | https://w.wiki/EghX |
| Purple | Mizunoryu, Badseed, Jacobolus | https://w.wiki/AGiB |
| Red | ThunderBrine | https://w.wiki/A9QR |
| White | Badseed | https://w.wiki/EghZ |
| Yellow | Badseed | https://w.wiki/Egha |

**Table S2.** Performance comparison between the automated charisma classification workflow (with a 5% color-proportion threshold) and expert classification across color classes. Each column reports the total number of hits (true positives), misses (false negatives), false alarms (false positives), and correct rejections (true negatives) in the charisma dataset. The percentage of correct hits (parentheses) are calculated as the number of hits divided by the total number of color targets (hits + misses) in the expert dataset. The percentage of correct rejections (parentheses) are calculated as the number of correct rejections divided by the total number of non-targets (correct rejections + false alarms) in the expert dataset.

| Color | Hits | Misses | Correct Rejections | False Alarms |
| --- | --- | --- | --- | --- |
| Black | 31 (100%) | 0 | 1 (100%) | 0 |
| Blue | 17 (63%) | 10 | 5 (100%) | 0 |
| Brown | 14 (82%) | 3 | 6 (40%) | 9 |
| Green | 19 (90%) | 2 | 10 (91%) | 1 |
| Grey | 3 (100%) | 0 | 13 (45%) | 16 |
| Orange | 0 (0%) | 2 | 27 (90%) | 3 |
| Purple | 0 (100%) | 0 | 32 (100%) | 0 |
| Red | 0 (0%) | 2 | 30 (100%) | 0 |
| White | 3 (33%) | 6 | 22 (96%) | 1 |
| Yellow | 9 (47%) | 10 | 13 (100%) | 0 |

**Table S3.** Performance comparison between the automated charisma classification workflow (with a 7.5% color-proportion threshold) and expert classification across color classes. Each column reports the total number of hits (true positives), misses (false negatives), false alarms (false positives), and correct rejections (true negatives) in the charisma dataset. The percentage of correct hits (parentheses) are calculated as the number of hits divided by the total number of color targets (hits + misses) in the expert dataset. The percentage of correct rejections (parentheses) are calculated as the number of correct rejections divided by the total number of non-targets (correct rejections + false alarms) in the expert dataset.

| Color | Hits | Misses | Correct Rejections | False Alarms |
| --- | --- | --- | --- | --- |
| Black | 30 (97%) | 1 | 1 (100%) | 0 |
| Blue | 15 (60%) | 10 | 7 (100%) | 0 |
| Brown | 13 (76%) | 4 | 7 (47%) | 8 |
| Green | 19 (86%) | 3 | 10 (100%) | 0 |
| Grey | 3 (100%) | 0 | 18 (62%) | 11 |
| Orange | 0 (0%) | 2 | 28 (93%) | 2 |
| Purple | 0 (100%) | 0 | 32 (100%) | 0 |
| Red | 0 (0%) | 2 | 30 (100%) | 0 |
| White | 3 (33%) | 6 | 23 (100%) | 0 |
| Yellow | 7 (37%) | 12 | 13 (100%) | 0 |

**Table S4.** Performance comparison between the automated charisma classification workflow (with a 10% color-proportion threshold) and expert classification across color classes. Each column reports the total number of hits (true positives), misses (false negatives), false alarms (false positives), and correct rejections (true negatives) in the charisma dataset. The percentage of correct hits (parentheses) are calculated as the number of hits divided by the total number of color targets (hits + misses) in the expert dataset. The percentage of correct rejections (parentheses) are calculated as the number of correct rejections divided by the total number of non-targets (correct rejections + false alarms) in the expert dataset.

| Color | Hits | Misses | Correct Rejections | False Alarms |
| --- | --- | --- | --- | --- |
| Black | 30 (97%) | 1 | 1 (100%) | 0 |
| Blue | 13 (52%) | 12 | 7 (100%) | 0 |
| Brown | 13 (76%) | 4 | 9 (60%) | 6 |
| Green | 17 (77%) | 5 | 10 (100%) | 0 |
| Grey | 3 (100%) | 0 | 20 (69%) | 9 |
| Orange | 0 (0%) | 2 | 29 (97%) | 1 |
| Purple | 0 (100%) | 0 | 32 (100%) | 0 |
| Red | 0 (0%) | 2 | 30 (100%) | 0 |
| White | 2 (22%) | 7 | 23 (100%) | 0 |
| Yellow | 5 (26%) | 14 | 13 (100%) | 0 |

